# Extensive splicing across the Saccharomyces cerevisiae genome

**DOI:** 10.1101/515163

**Authors:** Stephen M. Douglass, Calvin S. Leung, Tracy L. Johnson

## Abstract

Pre-mRNA splicing is vital for the proper function and regulation of eukaryotic gene expression. *Saccharomyces cerevisiae* has been used as a model organism for studies of RNA splicing because of the striking conservation of the spliceosome components and its catalytic activity. Nonetheless, there are relatively few annotated alternative splice forms, particularly when compared to higher eukaryotes. Here, we describe a method to combine large scale RNA sequencing data to accurately discover novel splice isoforms in *Saccharomyces cerevisiae*. Using our method, we find extensive evidence for novel splicing of annotated intron-containing genes as well as genes without previously annotated introns and splicing of transcripts that are antisense to annotated genes. By incorporating several mutant strains at varied temperatures, we find conditions which lead to differences in alternative splice form usage. Despite this, every class and category of alternative splicing we find in our datasets is found, often at lower frequency, in wildtype cells under normal growth conditions. Together, these findings show that there is widespread splicing in *Saccharomyces cerevisiae*, thus expanding our view of the regulatory potential of RNA splicing in yeast.

**Author Summary:** Pre-mRNA splicing is a fundamental step in eukaryotic gene expression. *Saccharomyces cerevisiae*, also known as brewer’s yeast, is a model organism for the study of pre-mRNA splicing in eukaryotes. Through the process of pre-mRNA splicing, a single gene is capable of encoding multiple mature mRNA products, but it is often difficult to identify the splice events that lead to these mRNA products. Here, we describe a method to accurately discover novel splice events in *Saccharomyces cerevisiae* and find evidence for extensive splicing in *Saccharomyces*. By utilizing a variety of strains and growth conditions, we are able to characterize many splice forms and correlate cellular conditions with prevalence of novel splice events.

## Introduction

Eukaryotic genes are composed of coding sequences termed exons interrupted by non-coding sequences called introns. Introns are removed from RNA by the large macromolecular complex known as the spliceosome through the process of RNA splicing. By selectively including different combinations of exons, a single gene can produce multiple RNA products. Pre-mRNA splicing is crucial for the proper expression of eukaryotic genes, and spliceosome components are highly conserved from yeast to mammals at the sequence, structure, and functional levels [1, 2].

Despite the high conservation between the spliceosomal components in yeast and mammals, *Saccharomyces cerevisiae* has a more compact genome with approximately 300 annotated intron-containing genes. Even with the relatively streamlined splicing landscape in *S. cerevisiae*, the prevalence of alternative splicing in this organism has not been well characterized. Several studies have found instances of alternative splicing in individual genes in *S. cerevisiae* [3, 4, 5], and there have been high-throughput methods to find novel splicing [6, 7, 8, 9, 10]. Most of these methods focus primarily or entirely on intron retention or exon skipping, while there has been little description of novel 5’ or 3’ splice site usage.

Here we describe a method for discovery of unannotated splice sites in *Saccharomyces cerevisiae* by RNA-seq. We tested the method with wildtype strains as well as strains and conditions that do not lead to direct changes in the splicing machinery itself, but that impact broad cellular conditions, RNA turnover, chromatin remodeling, and histone composition. Many of these strains and conditions help us observe changes in splicing that are not or are rarely detected in wildtype cells under normal growth conditions. We also analyze several strains that include *prp43-1*, a temperature sensitive mutation of the *prp43* gene, an RNA helicase directly involved in spliceosome disassembly to determine if modulating disassembly might affect our ability to detect aberrant splice site sampling. Interestingly, this mutation of *prp43* leads to a decrease in splicing of both annotated introns as well as novel splicing.

We compare our results with those of earlier studies and find good agreement, however over two thirds of the splice forms our method predicts, 676 total novel splice forms, have not been previously described. Within known intron-containing genes (ICGs), we find that novel isoforms generate longer introns in samples grown at higher temperature. We also find that deletion of *xrn1*, an evolutionarily-conserved 5’ to 3’ exonuclease, leads to an accumulation of novel splice isoforms that do not use either an annotated 5’ nor an annotated 3’ splice site, both within known ICGs and unannotated ICGs. While most of our novel splice forms use a known 5’ splice site and a novel 3’ splice site, we find that deletion of *ume6*, a component of a histone deacetylase complex that regulates early meiosis and that has been recently shown to have a regulatory role during mitosis [11], causes a dramatic increase in the use of novel 5’ splice sites within ICGs. We also find evidence for splicing of RNAs that are antisense to annotated transcripts. Together, our results indicate that there is significantly more splicing than previously thought. This suggests that the opportunity for splicing in the form of “latent” introns is a key feature of the yeast genome. Furthermore, changes in the activity of the gene expression machinery or the cells’ environment can significantly alter this rich splicing landscape.

## Results

### Alternative splicing is widespread in *Saccharomyces cerevisiae*

In order to explore the extent of splicing across the yeast transcriptome in a high confidence manner, we implemented a novel approach to allow us to leverage a large amount of RNA sequence data while imposing stringent filters (figure 1). Briefly, a large number of RNA-seq datasets are combined such that each novel splice form needs at least a single read that aligns without mismatches to the novel junction to pass the initial filter. This read cannot also align to known transcripts or the *Saccharomyces cerevisiae* genome, even with a large number of mismatches. Once a splice junction has been identified in this way, all reads are realigned to the newly discovered splice forms, annotated transcripts, and the genome to find the optimum alignment. Each novel splice form is then scored based on how well its 5’ splice site, 3’ splice site, and predicted branch point fit the consensus sequence for *Saccharomyces cerevisiae* splice signals. These scores are then used to compute p-values that represent how likely a splice product score of this strength is to occur by chance (figure S1).

**Figure 1.**
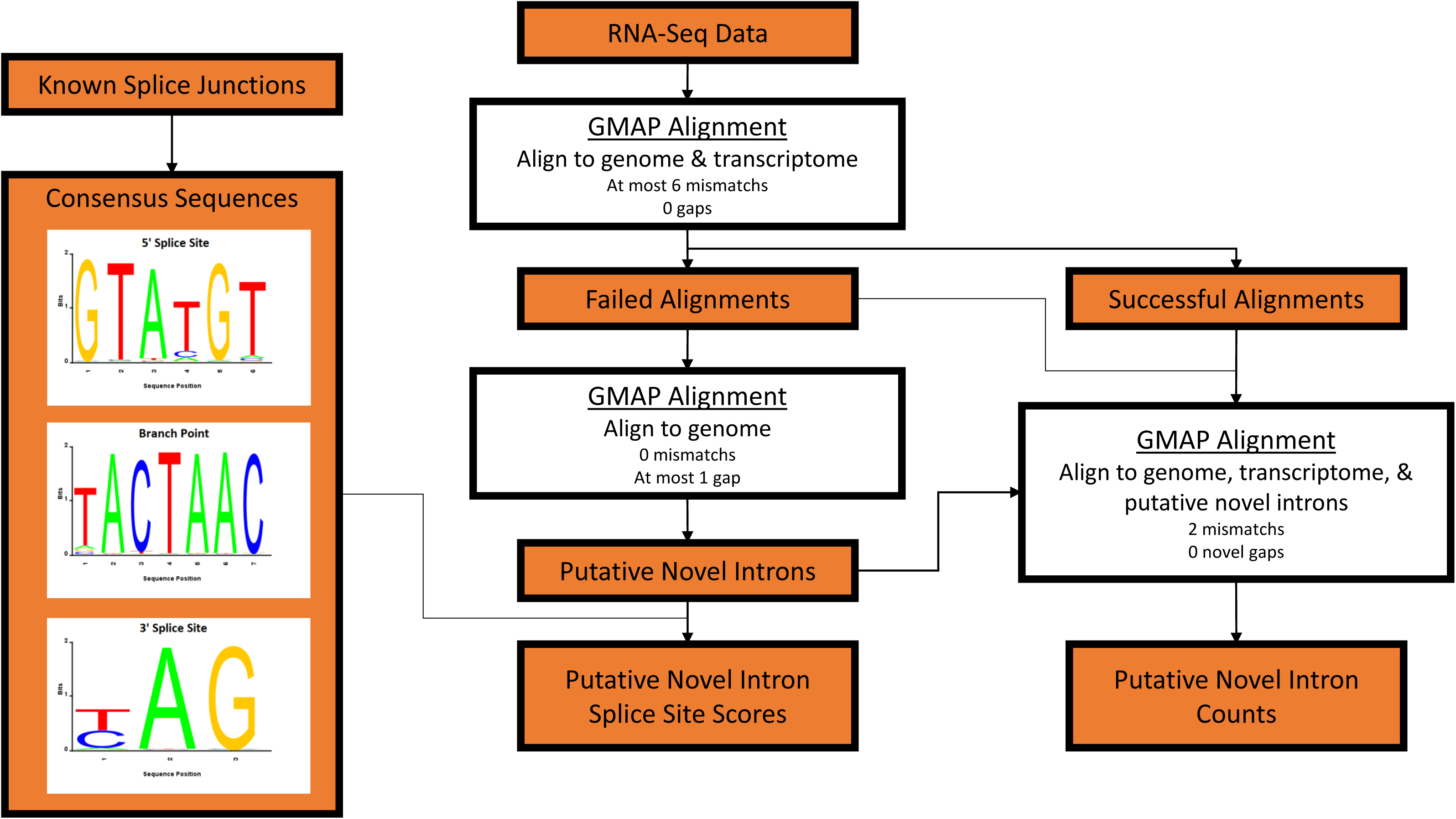
Workflow for discovery of novel splice forms. Multiple RNA sequence datasets are consolidated then aligned to the *Saccharomyces cerevisiae* genome and annotated transcripts liberally, allowing up to six mismatches but no gaps. Reads that fail to align are again aligned, but now allowing a single gap corresponding to a putative intron and no mismatches. This step defines all putative novel introns. Once again all reads are aligned to the *Saccharomyces cerevisiae* genome, known transcripts, and putative novel introns allowing up to two mismatches, which determines each putative novel intron’s read counts. Consensus sequences are determined from known *Saccharomyces cerevisiae* 5’ splice site, 3’ splice site, and branch point sequences, and each putative novel intron is scored based on sequence similarity to the consensus sequences.

To capture as many novel splice forms as possible, we incorporated datasets from a variety of strains and experimental conditions that are known or suspected to have an impact of splicing, some accessed from previous studies [12, 13] and some novel (Table S1). In addition to 4 wildtype samples grown at three different temperatures, these samples include 2 biological replicates each of two deletions in genes involved in decay, including *xrn1*, a 5’-3’ exonuclease, and *upf1*, required for efficient nonsense-mediated decay. We also include *htz1*Δ which encodes the histone variant H2A.Z and *swr1*Δ which is required to exchange H2A for H2A.Z in chromatin as well as double mutants *swr1*Δ *xrn1*Δ, *swr1*Δ *upf1*Δ, *htz1*Δ *xrn1*Δ, and *htz1*Δ *upf1*Δ. Cells that lack H2A.Z are found to have impaired splicing of intron-containing genes (ICGs), particularly genes that have suboptimal splice sites [13, 14]. We also analyze RNA-seq data from *snf2*Δ, which leads to an increase in use of non-canonical branch point and 5’ splice site sequences in annotated ICGs and *ume6*Δ, which derepresses genes implicated in meiosis in *Saccharomyces cerevisiae*, and the double mutants *upf1*Δ *snf2*Δ, *xrn1*Δ *snf2*Δ, and *ume6*Δ *snf2*Δ. Previous studies suggest that *snf2*Δ increases splicing by altering ribosomal protein gene expression and *ume6*Δ allows expression of genes that are usually repressed. We also include *set1*Δ and *set2*Δ, deletion mutations in histone methyltransferase genes. Finally, since we previously showed that the DEAH protein Prp43 contributes to disassembly of suboptimal spliceosomes using a *prp43* DAMP allele [13], we included a temperature sensitive mutant of *prp43, prp43-1*, as well as *set1*Δ *prp43-1* and *set2*Δ *prp43-1*. All *set1*Δ, *set2*Δ, and *prp43-1* strains contribute two samples to our workflow, one grown at 25° and one at 37°. Taken together, these datasets represent 29 samples across a variety of *Saccharomyces cerevisiae* strains and growth temperatures. By leveraging a large number of datasets, we are able to discover more novel splice sites and determine cellular conditions that lead to changes in alternative splice site usage.

| Strain | Read counts | Aligned read counts |
| --- | --- | --- |
| Wildtype | 20395478 | 14160763 |
| *xrn1Δ* | 36132564 | 25787231 |
| *upf1Δ* | 32005528 | 22034981 |
| *swr1Δ* | 38830226 | 21964031 |
| *htz1Δ* | 35969862 | 18508451 |
| *swr1Δ xrn1Δ* | 86744422 | 50807249 |
| *swr1Δ upf1Δ* | 11369368 | 6729660 |
| *htz1Δ xrn1Δ* | 52891892 | 35039812 |
| *htz1Δ upf1Δ* | 49460562 | 25424884 |
| Wildtype | 35059908 | 27722481 |
| *snf2Δ* | 34016283 | 25657587 |
| *upf1Δ* | 21328241 | 14465110 |
| *upf1Δ snf2Δ* | 31564582 | 23487340 |
| *xrn1Δ* | 29688304 | 20282088 |
| *xrn1Δ snf2Δ* | 34289534 | 27043467 |
| *ume6Δ* | 40071056 | 30161996 |
| ume6Δ snf2Δ | 22534282 | 17424922 |
| Wildtype 25° C | 126355512 | 97015734 |
| Wildtype 37° C | 140962504 | 82278728 |
| set1Δ 25° C | 120413012 | 75538896 |
| set1Δ 37° C | 150376422 | 84044314 |
| set2Δ 25° C | 115930956 | 61396568 |
| set2Δ 37° C | 105592626 | 63182213 |
| prp43-1 25° C | 113138574 | 78671916 |
| prp43-1 37° C | 93850396 | 54354046 |
| prp43-1 set1Δ 25° C | 180103858 | 107502423 |
| prp43-1 set1Δ 37° C | 116668118 | 70474357 |
| prp43-1 set2Δ 25° C | 129498126 | 93547450 |
| prp43-1 set2Δ 37° C | 107964066 | 68127119 |

Using our method, we discover evidence for 944 novel splice events across 408 transcripts with p < 0.05 (Table 1). Of these novel events, the majority are novel splice products found within known ICGs, using either the annotated 5’ splice site with a novel 3’ splice site (Table S2, n=537) or using the annotated 3’ splice site with a novel 5’ splice site (Table S2, n=129). Additionally, we found several cases of novel splicing that do not use any annotated splice sites within annotated ICGs (Table S2, n=22) and in unannotated ICGs (Table S2, n=198). We also find evidence for splice forms that are antisense to known transcripts (Table S2, n=50).

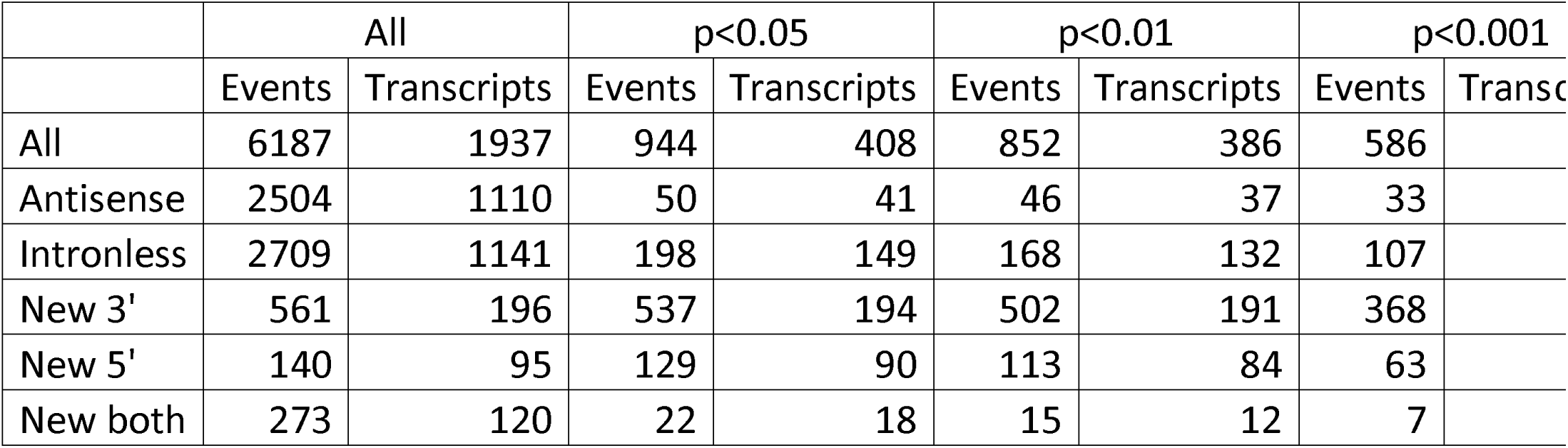

Many of the events we discover using our method are low abundance. About half of the total events are represented by fewer than five read counts across our datasets (figure S2A). However, these low abundance events are high confidence due to our approach’s stringent discovery protocol and p-value cutoff. Furthermore, these high confidence, low abundance novel splice events reveal splicing that is unlikely to be found in the high abundance data. Different classes of novel splicing are more prevalent in our low abundance novel splicing data. Specifically, novel splicing within annotated ICGs is more common in our high abundance data, while novel splicing within unannotated ICGs and antisense RNAs are more common in our low abundance data (figure S2C).

### Validation of novel splicing

In order to best validate our method by RT-PCR, we chose candidates with a variety of read counts and splice site scores representing each category of novel splicing we find, including novel 5’ splice site with annotated 3’ splice site, novel 3’ splice site with annotated 5’ splice site, both novel splice sites within an annotated ICG, novel splicing in an unannotated ICG, and splicing of transcripts that are antisense to annotated genes. In addition, we selected some genes which show only a single novel splice form and others with several. The results of our validation are shown in Figure 2 using oligonucleotides listed in table S3. *BIG1*, an integral membrane protein gene of the endoplasmic reticulum, shows a single novel splice event utilizing a novel 3’ splice site with an annotated 5’ splice site in our data with 19 total read counts spread roughly evenly across our 29 samples and a splice site score p-value of 0.0003. The novel splice event in *BIG1* is in-frame with the annotated form, and would be predicted to generate a protein product that has 6 additional amino acids. *SIM1*, a gene thought to participate in DNA replication, shows a total of four novel splice forms, each utilizing an annotated 3’ splice site and novel 5’ splice site with total read counts ranging from 2 to 35 and splice site p-values ranging from 0.018 to 0.0004. The intron in *SIM1* is located in the 5’ UTR, therefore the novel splice events would not be expected to yield a new protein product, although potential regulation in the 5’ UTR could be altered. *MCR1*, a gene involved in ergosterol biosynthesis in mitochondria, shows a single novel splice form utilizing both novel 5’ and 3’ splice sites within an annotated ICG with 49 total read counts, 9 of which are found across our *snf2*Δ samples, and a splice site p-value of 0.004. The annotated and novel introns for *MCR1* are both found in the 5’ UTR and are therefore unlikely to yield different protein products. However, we previously showed that changes in *SNF2* expression can affect splicing of others transcripts to alter mitochondrial function [15]. *SPF1*, which encodes an ion transporter of the ER membrane, is an unannotated ICG that shows five novel splice forms in our data with read counts ranging from 1 to 4 across all of our samples and p-value scores ranging from 0.004 to 6 × 10^-6^. Of the five novel splice forms found in *SPF1*, only one is in frame, which would produce a protein product that has 134 fewer amino acids. Finally, we find evidence for splicing in the antisense direction to *LEU4*, with a total of six splice forms with counts ranging from 1 to 33 across all samples and splice site p-values ranging from 0.0001 to 2.8 × 10^-6^. Not surprisingly, of the 48 sequence reads that derive from spliced reads antisense to *LEU4*, 28 come from samples which are deleted of *SET2*, a histone methyltransferase that has previously been implicated in suppressing antisense transcription [16]. Together, we validate each category of novel splicing we observe in our data and validate candidates ranging from a single read count to 49 read counts and from splice site p-values ranging from 0.018 to 2.8 × 10^-6^.

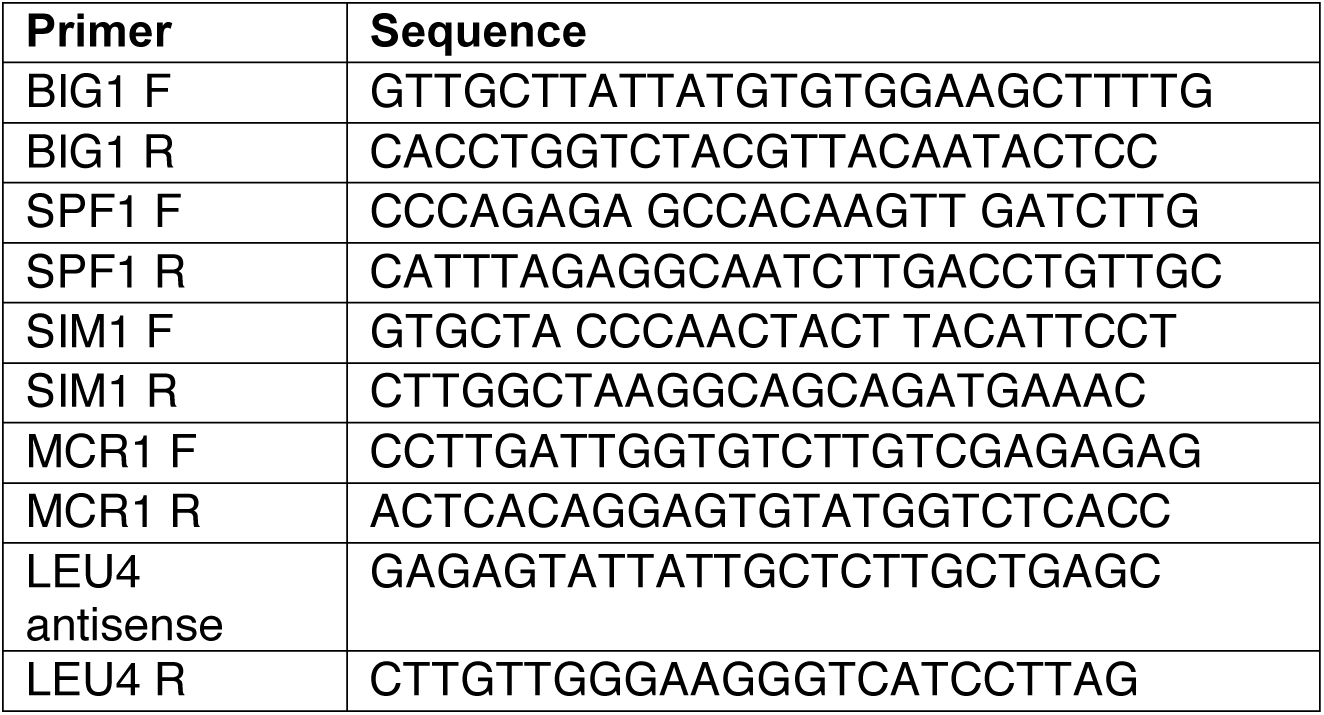

**Figure 2.**
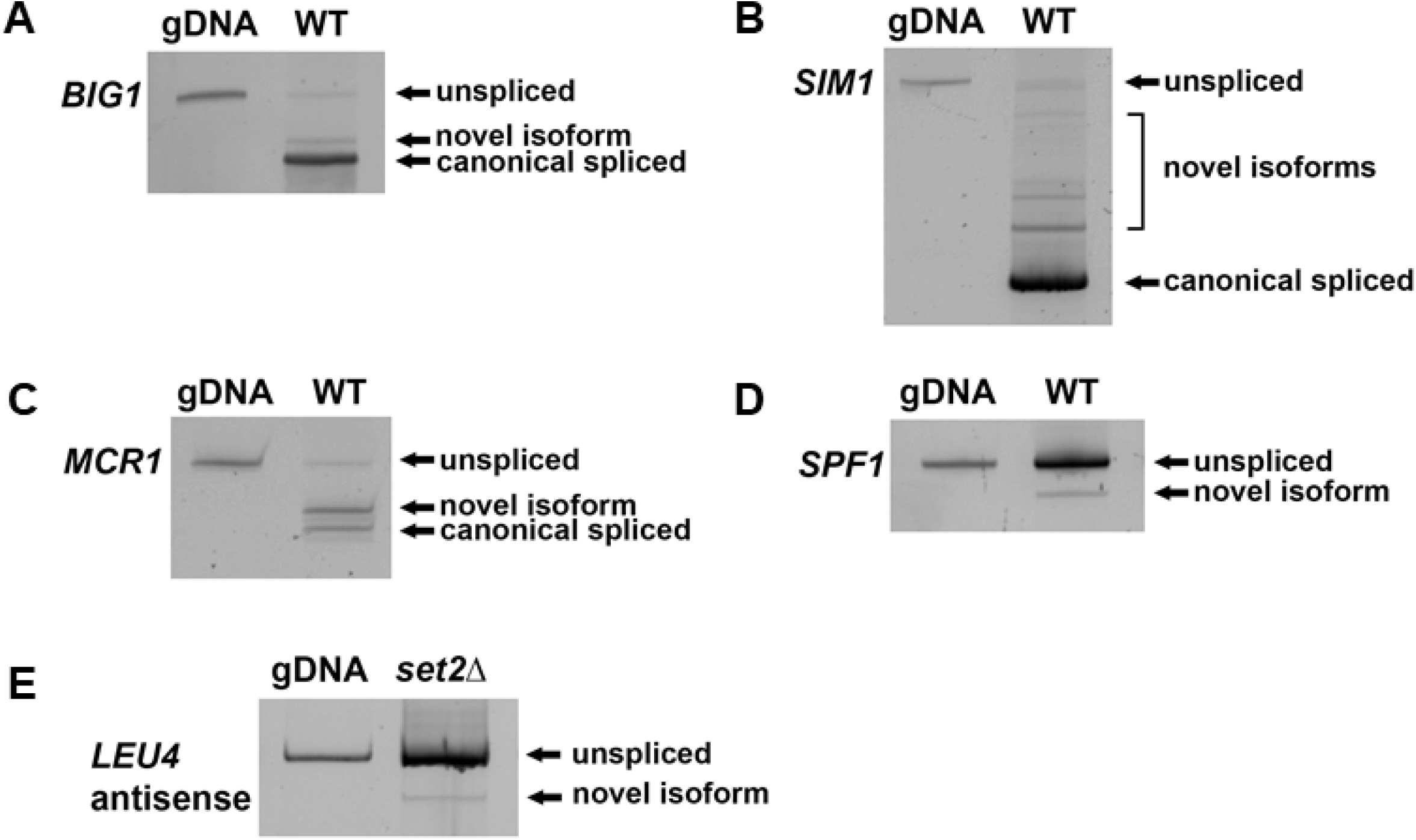
RT-PCR of representative novel splice isoforms. (A) *BIG1;* novel 3’ splice site within annotated ICG. (B) *SIM1*; 5’ UTR intron, novel 5’ splice site within annotated ICG. (C) *MCR1*; 5’ UTR intron, novel 5’ and 3’ splice sites within an annotated ICG. (D) *SPF1*; intron within an unannotated ICG. (E) *LEU4* antisense; antisense transcript of the *LEU4* gene containing an intron.

### Comparison with other sequence-based approaches for alternative splicing

Even though our method filters (1) putative splice forms with strong similarity to the *Saccharomyces cerevisiae* genome, (2) known transcripts, and (3) those found outside of genes, the splice sites identified by our method include many reported in recent studies. Of the 522 novel splice sites described in Kawashima et al. [6], 420 are discoverable by our method, 236 are found in our raw data and 189 pass our p < 0.05 filter. Of those described in Schreiber et al. [7], 248 out of 314 of the described splice events are discoverable by our method, 214 are in our data and 185 pass our p < 0.05 filter. Qin et al. [10] describe a method of identification of novel splice forms by lariat sequencing, a process which reveals 5’ splice sites and branch points but not 3’ splice sites. Of the 45 novel 5’ splice sites found in their work, 11 are present in our raw data and 9 pass our p < 0.05 filter with at least one corresponding 3’ splice site. Gould et al. [9] combined lariat sequencing with RNA-seq to identify both 5’ and 3’ splice sites along with novel branch point sequences. Of the 213 novel splice sites they report, 194 are discoverable by our method, 133 are found in our raw data, and 114 pass our p < 0.05 filter. The overlap between our work and these previous studies illustrates the power of our approach. None of the splice sites described by Qin et al. are found in any of the three other studies, while each of Kawashima et al., Gould et al., and Schreiber et al. have greater overlap with our work than with one another (figure 3). Taken together, of the 944 splice junctions predicted with p < 0.05 by our method, 268 have been previously described and 676 are novel.

**Figure 3.**
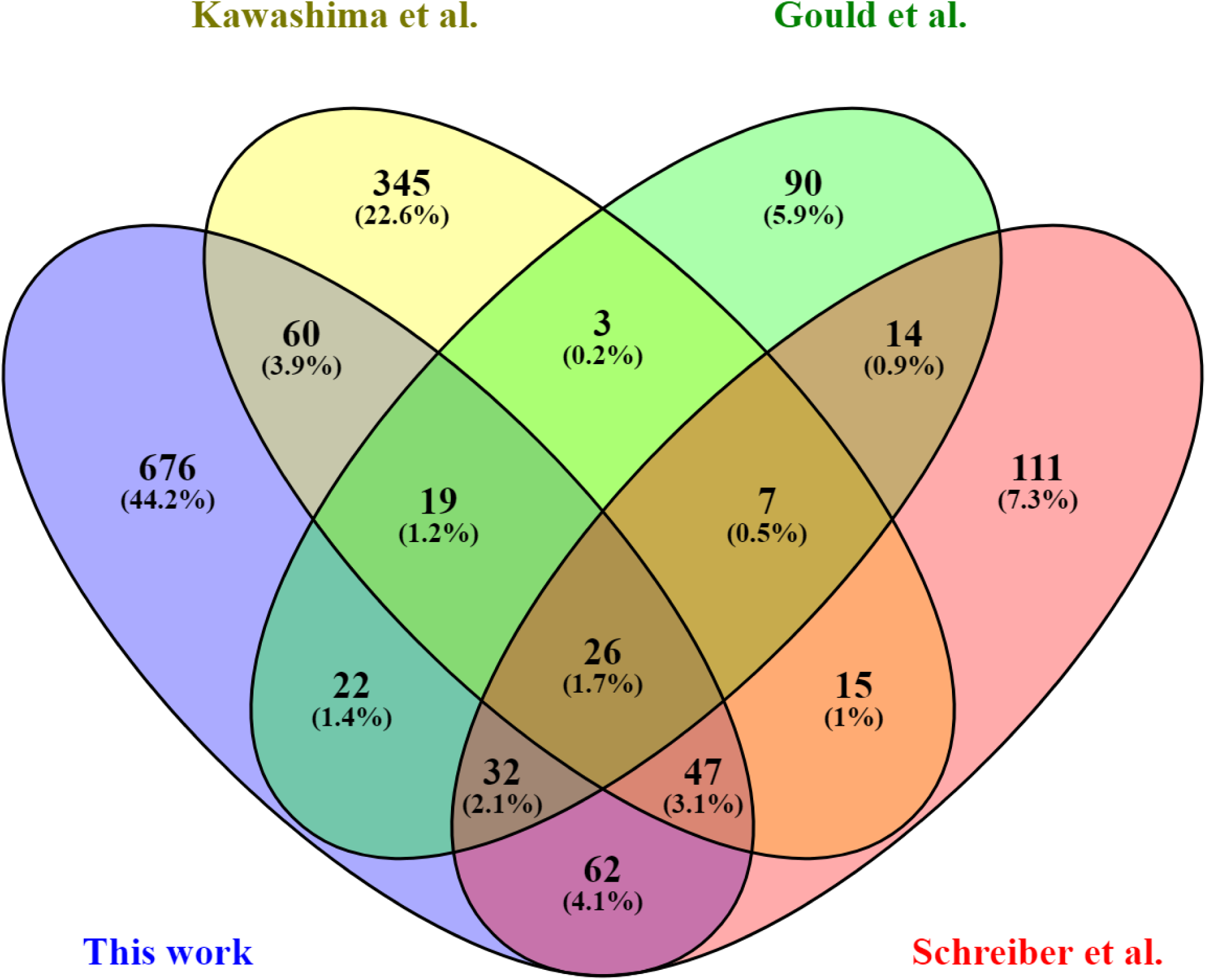
Agreement with previous studies. Venn diagram showing the overlap between the novel splice forms discovered here and those described previously.

### *Saccharomyces cerevisiae* contains antisense transcripts that undergo splicing

In addition to discovering novel splice sites within genes already known to contain introns and unannotated ICGs, our method allows us to discover splicing that is antisense to annotated transcripts, primarily in the degradation mutant strains *xrn1*Δ and *upf1*Δ and the histone methyltransferase mutant *set2*Δ. Previous studies report that Set2 suppresses antisense transcription [16]. We find evidence for 50 antisense novel splice events spread across 41 transcripts that pass our statistical criteria. Of these 41 transcripts, 37 show a single novel splice event, two transcripts have two unique novel splice events, one transcript has 3 separate events, and a single transcript, which is antisense to the *LEU4* gene, has 6 unique novel splice events. While most of the antisense splice events in our data have low read counts, the *LEU4* isoforms together account for 51 reads across 11 samples, including wildtype, *set2*Δ, and *set2*Δ *prp43-1* at both 25° and 37°, *set1*Δ and *set1*Δ *prp43-1* at 37° only, and *xrn1*Δ, *swr1*Δ *upf1*Δ, and *upf1*Δ *snf2*Δ. Together, *LEU4* antisense splicing represents over 20% of our total antisense read counts across all samples. These six isoforms arise from 3 different 5’ splice sites and 4 3’ splice sites. 5 out of 6 of these isoforms generate similar mature mRNAs with an intron in the size range from 115 nucleotides to 129 nucleotides and are therefore indistinguishable by RT-PCR (figure 2E). The remaining form is generated from a unique 5’ splice and 3’ splice and causes an intron of 464 nucleotides, and is low abundance in our RNA sequence data, with only a single read count in a single sample, *set2*Δ at 37°.

In unannotated ICGs, we find 149 transcripts that show novel splicing in the sense direction and 37 that undergo novel splicing in the antisense direction. Interestingly, the number of transcripts that show novel splicing in both the sense and antisense direction is just one, *DJP1*. If a set of 149 genes and a set of 37 genes are each randomly chosen from all *Saccharomyces cerevisiae* transcripts, the expected value of the overlap is a single transcript, suggesting that within unannotated ICGs, presence of novel splicing in the sense direction does not significantly impact the chances of novel spicing of an antisense transcript or vice versa. Within annotated ICGs, 220 transcripts undergo novel splicing in the sense direction and 4 have novel splicing in the antisense direction, indicating no correlation between the splicing of annotated intron-containing genes and their corresponding antisense transcripts.

### Prp43 is required for efficient splicing of annotated and novel introns

Several of the strains in our analysis include *prp43-1*, a temperature sensitive mutation that is viable at 25° C but not at 37° C. Prp43 is an RNA helicase that has a vital role in spliceosome disassembly and is required for efficient mRNA splicing in *Saccharomyces cerevisiae*. *PRP43* has also been implicated in ribosome biogenesis [17], and we previously showed that decreasing levels of *PRP43* using a DAMP allele can suppress splice defects [13]. To characterize the consequences of the *prp43-1* mutation in splicing, we compared the *prp43-1* strain to a wildtype strain, a *set1*Δ *prp43-1* strain to a *set1*Δ strain, and a *set2*Δ *prp43-1* strain to a *set2*Δ strain at both 25° and 37° C. As expected from its role in splicing, *prp43-1* shows a decrease in the splicing of annotated introns in *Saccharomyces cerevisiae* (figure 4A). Interestingly, despite earlier findings that reducing levels of wildtype Prp43 can suppress splice defects and promote splicing of weak introns [13], we find that strains with *prp43-1* show less novel splicing than their counterparts with wildtype *PRP43* (figure 4B).

**Figure 4.**
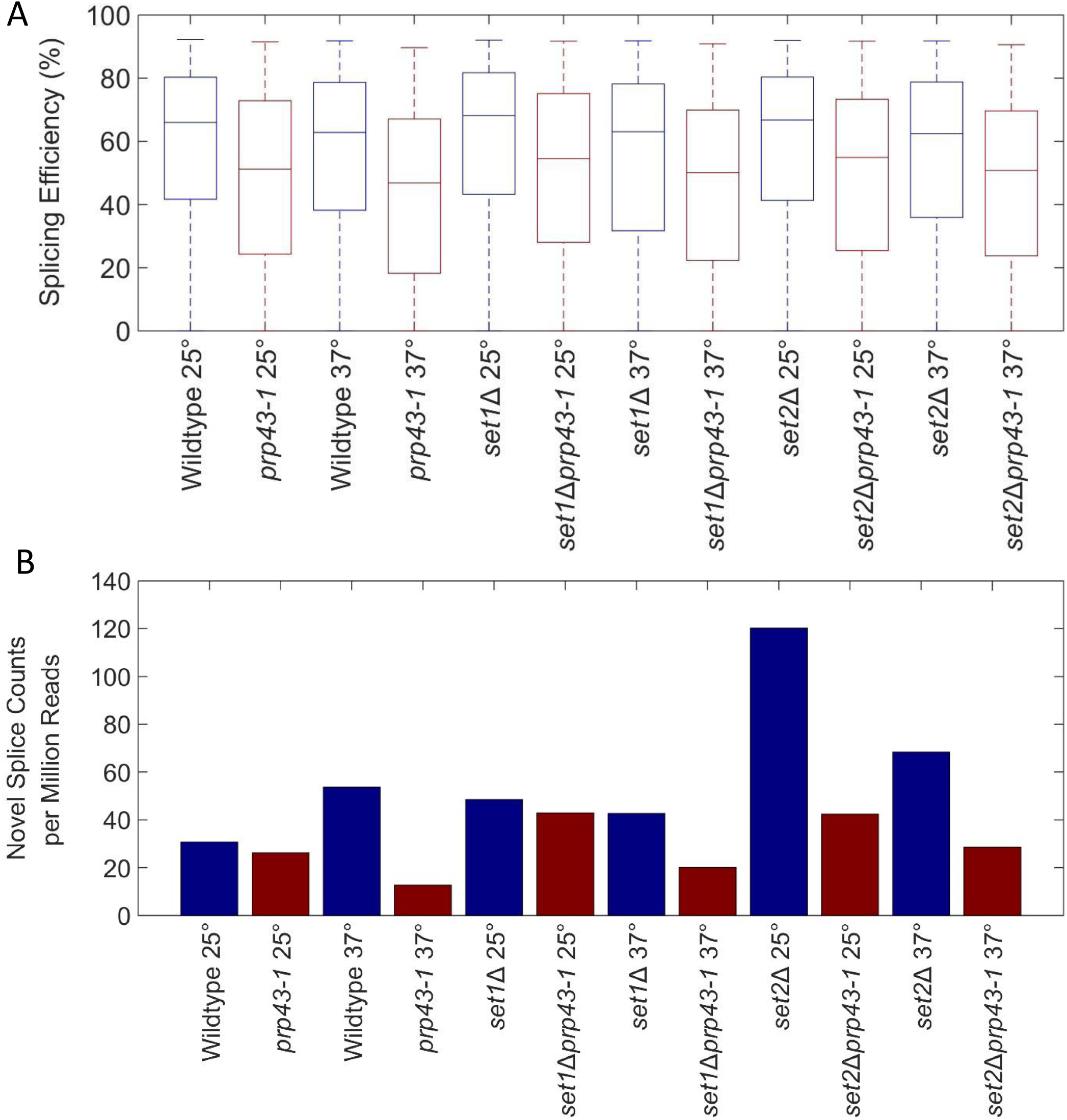
*prp43-1* leads to a decrease in splicing. (A) Strains with the mutated *prp43-1* gene show a decrease in splicing efficiency of annotated introns relative to strains with wildtype *PRP43*. (B) Evidence of novel splicing is lower in strains with *prp43-1* versus wildtype *PRP43*.

### Elevated temperature favors novel introns that are longer than their annotated introns

Across our datasets, we detect fewer novel splice events in high temperature samples than would be expected by sequence depth alone. This is unsurprising given that many of our lower temperature samples are mutants that lead to accumulation of normally degraded products, such as *xrn1*Δ and *upf1*Δ. While the total number of novel splice events is underrepresented at high temperature, the novel splice products generated differ in intron length. When cells are grown at 37° C, novel splicing within annotated ICGs tends to favor intron sizes that are larger than novel splice forms from cells that are grown at 25° or 30° C (figure 5A). This can be explained by effects observed in our most common class of novel splice events, those that use an annotated 5’ splice site with a novel 3’ splice site. At higher temperature, these novel 3’ splice sites tend to be further downstream than the annotated splice sites. We find that most of the temperature-enriched splice events are more commonly found in our samples with wildtype *prp43*, since *prp43-1* decreases both annotated and novel splicing (figure 5B). These results are consistent with work that finds that intron structure can function to control alternative splicing in yeast [18]. Nonetheless, our data reveal a number of previously unreported events.

**Figure 5.**
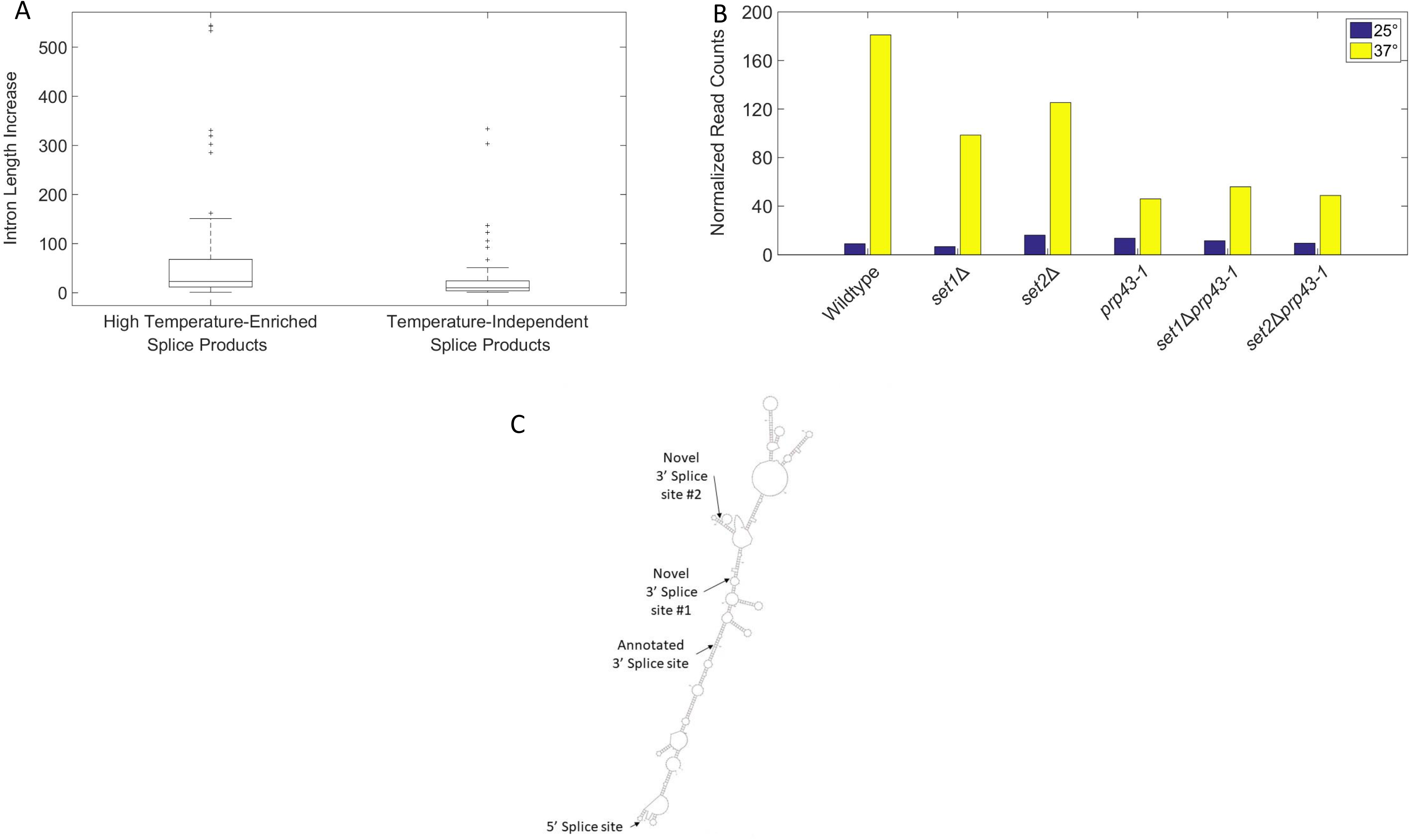
Elevated temperature leads to longer introns. (A) Boxplot of temperature-enriched splice forms versus temperature-independent splice forms shows higher temperature leads to an increase in intron size. Temperature-enriched splice forms have much higher counts at higher temperature, but are still present at lower temperature. (B) Shifting *prp43-1* to higher temperature causes a dramatic decrease in these temperature-enriched splice forms. (C) Predicted structure of TMH18 with annotated and novel splice sites labeled.

As an example, *TMH18*, a mitochondrial membrane protein gene, has an annotated intron that is 96 nucleotides long and two common novel splice isoforms discovered by our method, both of which generate longer introns. The isoform that is highly enriched in high temperature samples contains a 161 nucleotide intron, while the isoform that is not favored at high temperature contains a 128 nucleotide intron. We speculate that the increase in temperature destabilizes the secondary structure of some pre-mRNA molecules to allow access to normally inaccessible splice sites. This may lead to an increase in use of distant novel splice sites. To confirm this, we analyzed the predicted pre-mRNA secondary structure using MFOLD [19]. The predicted secondary structure with the most favorable free energy shows that the annotated 5’ splice site, the annotated 3’ splice site, and the novel 3’ splice site that is not favored at high temperature are all readily accessible, while the novel 3’ splice site that is favored only at high temperature is not (figure 5C).

Interestingly, these distant splice sites may be under evolutionary pressure to be unable to code for protein, as 74/84 (88%) of these temperature-enriched splice sites contain premature termination codons. This is similar to the fraction of PTC-generating events found in the totality of our novel splicing events in annotated intron-containing genes, with 572/636 (90%) containing premature termination codons. This agrees with results described by Kawashima et al. [6] that find that stress conditions, including heat shock, cause an increase in non-productive novel splice usage. Together these data suggest that alternative splicing may be a mechanism for reconfiguring the transcriptome in response to stress.

### Xrn1 deletion increases splice forms that do not use annotated splice sites

Novel splicing is found most commonly in strains in which *xrn1* has been deleted. These strains account for approximately 9% of our total sequence depth but 19% of our total novel splice counts. Interestingly, *xrn1* deletion mutants are particularly enriched in novel splice forms that do not use any annotated splice sites. Roughly 18% of novel splice form counts that use either an annotated 3’ or 5’ splice site derive from an *xrn1* deletion mutant, while 31% of those that utilize two novel splices are found in an *xrn1* deletion strain. Two examples of unannotated ICGs impacted strongly by *xrn1* deletion are *AGC1* and *MRM2*, both involved in mitochondrial function. Interestingly, these genes’ expression increases in *xrn1*Δ, and they each have their highest RPKM values in the five samples in which *xrn1* is deleted [12, 13]. The 50 unannotated ICGs that are most enriched in our *xrn1*Δ strains only have a single GO term in common, “intracellular membrane-bounded organelle,” further suggesting an impact on transcripts important for mitochondrial function. Interestingly, a 2012 study found that Xrn1 is critical for the translation of genes necessary for mitochondrial function in *Saccharomyces cerevisiae* [20]. Together, this suggests a role for Xrn1 in the regulation of alternative splice products of mitochondrial transcripts.

### Ume6Δ-derived increase in IC-RPG expression leads to increase in IC-RPG alternative splicing?

Our *ume6*Δ strain is highly enriched in novel splice events which utilize an annotated 3’ splice site with a novel 5’ splice site. Of the newly discovered splice sites which utilize a novel 5’ splice site and a canonical 3’ splice site, 59% are in intron-containing ribosomal protein genes (IC-RPGs). However, of the events that are found disproportionately more frequently in *ume6*Δ, 70% are in IC-RPGs. Our RNA sequence data suggests that IC-RPG expression increases slightly in *ume6*Δ. Novel splicing in IC-RPGs increases by 70%, and 5’ novel IC-RPG splicing increases by 91% in *ume6*Δ-containing samples relative to samples with wildtype ume6.

Previous studies have shown that Ume6 is degraded under meiotic conditions [21]. This raises the interesting possibility that genetic manipulations that remove Ume6 may lead to changes in the unannotated splicing landscape that are related to what occurs under meiotic conditions. Ongoing experiments will test this model.

## Discussion

In this study we show that there are many rarely utilized splice sites in *Saccharomyces cerevisiae*. Our methodology is capable of discovering many novel splice forms by utilizing a large number of RNA-seq datasets. We are able to do this in a robust, high-confidence manner by excluding reads that can be explained by sequencing error or genomic DNA contamination and filtering based on existing splice site census sequence data. Overall, we find a strong preference for novel splicing using a known 5’ splice site and a novel 3’ splice site within annotated ICGs, which represent over half of our statistically significant novel splice products. However, due to our high total sequencing depth and variety of strains and experimental conditions we are still able to find a relatively large number of novel splice forms that utilize known 3’ splice sites with novel 5’ splice sites, those that utilize novel 3’ and 5’ splice sites within annotated ICGs, those that utilize two novel splice sites within unannotated ICGs, and splicing of transcripts that are antisense to annotated genes. The large number of novel splice events that we discover allow us to correlate changes in splice site preferences with different mutant strains and experimental conditions. While it has been tempting to call splicing events that have not been previously annotated as “errors,” we believe that these data actually reveal the remarkable substrate flexibility of an evolutionarily conserved enzyme. Moreover, it stands to reason that if splicing is to contribute to adaptation and, ultimately, evolution of multicellular organisms, then an array of sequences, not simply those that match a strong consensus, need to be recognized and spliced. The results described here provide a window into this sequence landscape.

In the adjoining manuscript [22], the authors find 229 “protointrons” – rapidly evolving, inefficiently spliced introns. Of these, we define 60 as novel introns with an additional 9 found in our RNA sequence data but filtered out due to low splice site scores. The limited overlap between the methods highlights that neither of our methods has reached saturation. Furthermore, the 10 strains in our analysis with the greatest normalized overlap with Talkish et al. are the 10 strains that include either *xrn1*Δ or *upf1*Δ, suggesting that overlap between the methods is more driven by stabilization of protointron-containing splice products than biological similarity between “hungry spliceosome” conditions and the strains used here. Talkish et al. confirm by RT-PCR several splice events that would be removed from our analysis due to poor splice signals, indicating that our method’s stringency likely eliminates many true splice products. The approach used here scores putative novel splice sites by learning from annotated splice sites, so as additional protointrons with unusual splice sites are discovered and validated, our method’s ability to discover these forms will increase.

The analysis of several mutant strains allows insights into splice site selection in *Saccharomyces cerevisiae*. We find general trends in our data as well as specific effects for particular strains or conditions. For example, elevated temperature leads to an increase in novel splice form intron length, *xrn1*Δ leads to a large increase in novel splice forms that do not use an annotated splice site, and *ume6*Δ leads to an increase in novel splicing of intron-containing ribosomal protein genes.

Despite the fact that analyzing many mutant strains can increase our understanding of splice site selection, every new category and class of splice form that we find is observed in our wildtype data. While *xrn1*Δ dramatically increases the number of splice products that arise from use of two novel splice sites, we see many examples of splice products that use two novel splice sites in our wildtype samples. This holds true for elevated temperature, *ume6*Δ, and *set2*Δ, as well as splicing of transcripts that are antisense to known transcripts, splicing that uses a single annotated splice site, and splicing in unannotated ICGs. Together, this indicates that while mutant strains are useful for correlating genetic changes with splicing changes, even wildtype *Saccharomyces cerevisiae* in normal conditions are capable of producing a large variety of splice products. Taken together, these findings illustrate the diversity and depth of splicing in *Saccharomyces cerevisiae*, and also show the presence of latent introns that are found across the genome.

## Methods

### Public datasets

*Saccharomyces cerevisiae* wildtype, *xrn1*Δ, *upf1*Δ, *swr1*Δ, *htz1*Δ, *swr1*Δ *xrn1*Δ, *swr1*Δ *upf1*Δ, *htz1*Δ *xrn1*Δ, and *htz1*Δ *upf1*Δ strain RNA-seq data were download from GEO (accession number GSE97416). Additional wildtype, *upf1*Δ, and *xrn1*Δ RNA-seq data as well as *snf2*Δ, *ume6*Δ, *upf1*Δ *snf2*Δ, *xrn1*Δ *snf2*Δ, and *ume6*Δ *snf2*Δ strain RNA-seq data were also downloaded from GEO (accession number GSE94404).

### Yeast culture and Sequencing

Wildtype, *set1*Δ, *set2*Δ, *prp43-1, set1*Δ *prp43-1*, and *set2*Δ *prp43-1* strains were grown at 25°C to OD 0.3. Then, cultures were equally split, half remaining at 25°C and half shifting to 37°C for four hours. 10 mL of cells were pelleted from each sample and RNA extraction was performed. Then, 20 μg total RNA per sample was treated with DNAse I (Roche) and depleted of rRNA with the Ribo-Zero Gold rRNA Removal Kit (Illumina). RNA-seq libraries were prepared using the Illumina TruSeq v2 RNA Kit. 50 base pair paired-end reads were generated on an Illumina HiSeq 4000.

### Sequence alignment

RNA sequence data were combined and aligned to the *Saccharomyces cerevisiae* SacCer3 genome reference and Ares Lab Yeast Intron Database Version 3 [23] in a single step using STAR [24] allowing up to six mismatches and no unannotated gaps. Sequence reads that fail to align in this step are then aligned to the SacCer3 genome again, this time allowing no mismatches, a single gap, at most one alignment locus, and at least 15 nucleotide overhang on each end of the alignment. Successful alignments in this step define putative novel introns. In a final alignment step, all reads are aligned to the SacCer3 genome, the Ares Intron Database, and newly defined putative novel introns in one step, allowing at most 1 alignment locus with up to 2 mismatches. Counts for novel splice forms are based on those derived from this alignment step.

### Splice site scoring

5’ splice site, 3’ splice site, and branch point scores were generated based on how closely each splicing signal matches the consensus sequence for that signal. For each position within a splice site, the total number of adenine, cytosine, guanine, and thymine bases present at that position in annotated splice signals was determined based on the Ares Lab Intron Database, and then the proportion of each nucleotide at that position was found by dividing by the total number of annotated splice products. The score for each splice site was calculated as follows:

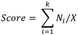

Where i is the position within the splice signal, k is the number of positions in the splice signal, N_i_ is the count of that specific nucleotide at that position, and X is the total counts for the splice signal. To score each putative novel splice form, the 5’ and 3’ splice sites are determined from RNA-seq data, while branch point sequences were chosen by selecting the highest scoring possible branch point sequence within a maximum distance of 200 nucleotides from the 3’ splice site. The score for each putative novel splice site is simply the product of its 5’ splice site score, its 3’ splice site score, and its branch point score. Putative novel splice sites were considered antisense if the score in the antisense direction is higher than the score in the sense direction.

### Statistical analysis of splice sites

P-values for putative novel splice sites were generated by comparing the potential splice site to all possible combinations of 5’ splice sites, 3’ splice sites, and branch points. Based on a six nucleotide 5’ splice site, a three nucleotide 3’ splice site, and a seven nucleotide branch point, there are 4,294,967,296 possible combinations of splice signals that can be described. To convert our putative novel splice form scores to p-values we compared the score to the scores of all possible combinations of splice signals. The raw p-value is the fraction of these combinations that score equal to or better than the putative novel form, which also represents the chance of seeing a splice score as good or better than this by chance. We then correct the raw p-values for multiple hypothesis testing using the Bonferroni correction by multiplying each raw p-value by the number of tests conducted, which is the total number of putative novel splice sites times two, since each splice site score is calculated in both the sense and antisense direction. All adjusted p-values greater than one are then set to one.

### Splicing Efficiency Calculation

To calculate the splicing efficiency of annotated splice sites, reads were aligned with STAR [24] allowing only a single alignment locus, only annotated splice sites, and at most two mismatches. Aligned reads within ICGs were categorized as exonic, spliced, or unspliced. We generated normalized spliced and unspliced read counts by dividing the raw counts in each category by the number of possible alignments that can fall into that category. This equates to read length minus one for spliced reads and the intron length plus the read length minus one for unspliced reads. Splicing efficiency is then computed as normalized spliced counts divided by the sum of normalized spliced and normalized unspliced counts.

### RNA Folding

To find the optimum secondary structure for TMH18, we used the MFOLD web server with the pre-mRNA sequence accessed from the Saccharomyces Genome Database (https://www.yeastgenome.org/) and default MFOLD parameters [19; http://unafold.rna.albany.edu/?q=mfold]. The optimum secondary structure was visualized using MFOLD.

### RNA isolation and RT-PCR

RNA was isolated from log phase yeast by hot phenol:chloroform:isoamyl alcohol (PCA) extraction with SDS. The RNA was then precipitated with ethanol. 20 µg of total RNA was DNase-treated with 30 U DNase 1 (Roche) for 1 hour at 25°C. 1 µg of DNase-treated RNA was used for cDNA synthesis. cDNA synthesis was performed using the Maxima First Strand cDNA Synthesis Kit (ThermoFisher). 1 µL of cDNA was used for the Taq PCR reaction using gene-specific primers to analyze splicing.

### Data Visualization

Venn diagrams to view the overlap between this and previous works were generated using Venny 2.1.0 [27]. Box plots and bar graphs were generated using the MATLAB functions boxplot and bar, respectively.

### Accession numbers

Data generated in this study is available under GEO accession number GSE120497.

## Acknowledgements

We thank the Dr. Tracy Johnson lab (UCLA) for comments and suggestions to improve the manuscript and Manuel Ares, Jr. for generously sharing unpublished data.

---

**Figure S1.**
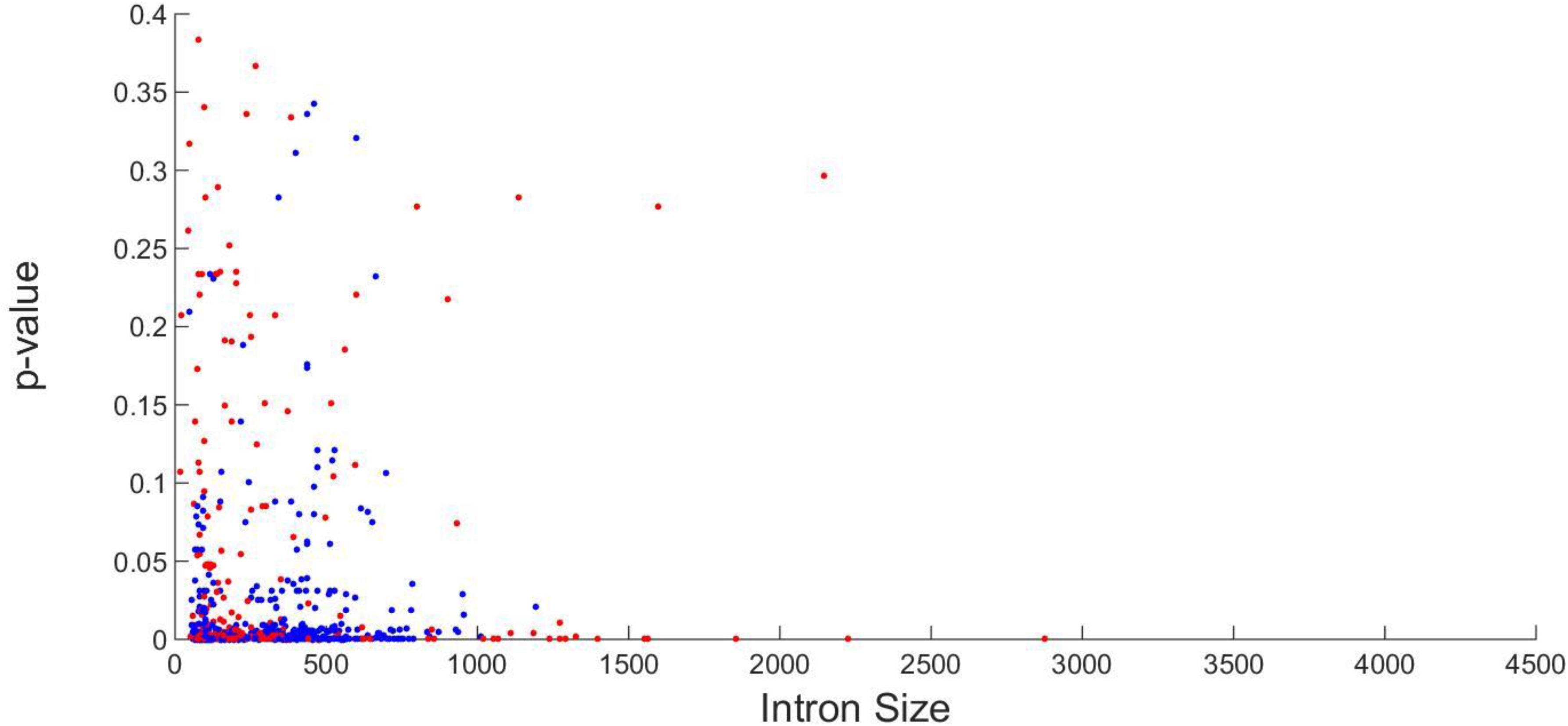
Putative novel splice site scores. Each putative novel splice site was scored by similarity to known *Saccharomyces cerevisiae* splice signals. The splice site score was plotted against putative intron length for novel splice products within annotated ICGs (blue) and unannotated ICGs (red).

**Figure S2.**
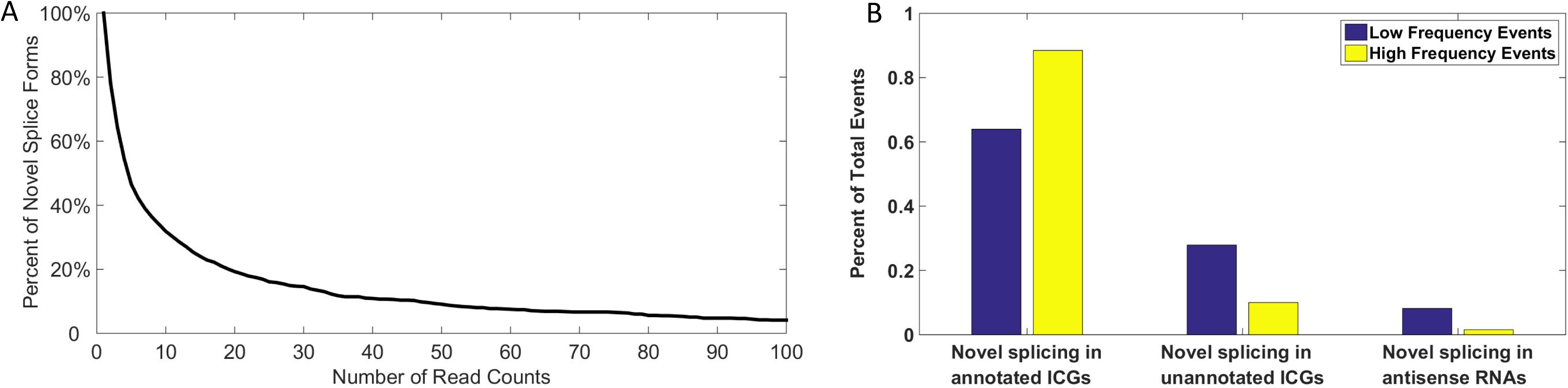
Summary of novel splice products. (A) Over half of the data have fewer than 6 counts across all 29 of our datasets. (C) Novel splicing outside of annotated intron-containing genes and splicing of RNAs that are antisense to known transcripts is observed.

